# Temporal niche pursuit in a simulated evolution of sleep/wake patterns

**DOI:** 10.64898/2026.05.24.726501

**Authors:** Krutika Ambani, John A. Lesku, Roelof A. Hut, Charles L. Nunn, Andrew J. K. Phillips

**Affiliations:** School of Psychological Sciences, Turner Institute for Brain and Mental Health, Monash University, Clayton, VIC 3800, Australia; School of Agriculture, Biomedicine and Environment, La Trobe University, Melbourne, VIC 3086, Australia; Chronobiology Unit, Groningen Institute for Evolutionary Life Sciences, University of Groningen, 9747 AG Groningen, The Netherlands; Department of Evolutionary Anthropology, Duke University, Durham, NC 27708, USA; Duke Global Health Institute, Duke University, Durham, NC 27708, USA; Flinders Health and Medical Research Institute (Sleep Health), Flinders University, Bedford Park SA 5042, Australia

## Abstract

It was long believed early mammals were nocturnal to avoid interactions with day-active dinosaurs. However, recent evidence indicates many dinosaurs were likely nocturnal, suggesting more complex coevolutionary dynamics prevailed. We simulated coevolution of sleep in a general predator/prey system, using a physiological model. We discovered ‘temporal niche pursuit’ cycles across evolutionary timescales: prey repeatedly escaping into a novel temporal niche, with predators subsequently invading that niche. We characterized multiple oscillatory patterns for pursuit, involving distinct genetic and phenotypic mechanisms. A low-dimensional model recapitulated the dynamics of the physiological model. These findings reveal rich dynamical processes underlying selection of temporal niche.

## I. INTRODUCTION

Evolutionarily, sleep is a highly conserved behavior, emerging before the brain [1,2]. Sleep phenotypes differ markedly between species, for example, humans sleep in consolidated nightly blocks, whereas mice sleep in multiple bouts, generally in the day. Temporal niche refers to the times of day a species is awake and capable of performing crucial behaviors, including feeding and mating [3]. Differences in temporal niche between species enable resource partitioning with competitors [4],[5],[6] and can reduce risk of predation [7]. However, the evolutionary drivers of divergent temporal niches between closely related species [8],[9] remain poorly understood.

The ‘nocturnal-bottleneck hypothesis’ proposed that early mammals evolved the nocturnal phenotype to minimize interactions with diurnal dinosaurs, who were thought to be cold-blooded and day-active [10]. Under this hypothesis, mammals expanded into other temporal niches only after the Cretaceous–Paleogene extinction event 66 million years ago [11]. However, eye structures of fossil remains suggest many predatory dinosaurs had already invaded the nocturnal niche during the Mesozoic period (252-66 million years ago) [12]. Furthermore, dinosaurs seemingly evolved thermoregulation supportive of night-time activity [13],[14],[15] and the nocturnal phenotype evolved multiple times among proto-mammals [16]. Evolution of temporal niche occupation may thus be more complex than previously assumed.

To understand the general dynamics of sleep evolution in predator-prey systems, we adapted the standard, physiologically based Phillips-Robinson model of sleep to represent agents (individuals) within these populations. This model has been applied to a wide range of questions [17–20], informing the physiological basis for inter-species differences in sleep. We then explored whether key dynamics of the physiological model of sleep evolution could be recapitulated by a simple low-dimensional model.

## II. PHYSIOLOGICAL MODEL OF PREDATOR-PREY SLEEP EVOLUTION

The Phillips-Robinson model was used to generate sleep/wake time series for each simulated agent [19]. This ordinary differential equation model represents neural circuitry that control sleep/wake states; the sleep homeostatic process, *H*, generating increasing sleep pressure during time awake; and the circadian pacemaker (variables *x, x*_*c*_) and its stimulation by retinal photoreceptors (variable *n*). For rapid computation, a Hard Switch reduction was applied [21] (parameter values in Table S1).

Each agent was characterized by its “genome” and “phenotype”. The genome is defined by 4 parameter values, which could vary within meaningful physiological ranges, with 100 evenly spaced alleles:(i) s eep drive offset, *D*_*O*_ (-30 to 13 mV), which controls sleep duration (higher = more sleep); (ii) the sleep homeostatic time constant, *x* (5 to 45 h), which governs sleep cycling (lower = more rapid sleep/wake cycling); (iii) the diurnality index, *a* (-1 to 1), which modulates the circadian drive (positive = diurnal; negative = nocturnal); and (iv) the intrinsic circadian period, *τ* (22 to 26 h), which controls the phase of circadian entrainment (higher = later phase).

For each agent, a 13-day time series was generated (initial conditions: *H*= 16, *x* = — 0.7, *x*_*c*_ = 0, *n* = 0.03, state=wake), omitting the first 10 days to reduce transients. The phenotype is defined as the 1×7201 time series (0.01-h epochs), denoted *R*_*g*_ for agent *g*, with values of 1 (wake) or 0 (sleep). A 10-minute transition interval (scored as 0.5) was added following each sleep/wake transition to account for minimum time in a state required for physiological utility (due to sleep inertia after awakening, and sleep initiation across brain regions [22]).

Each evolutionary simulation consisted of 500 non-overlapping generations, with 100 agents per population. Light was 1000 lux from 6:00 to 18:00, with 0 lux otherwise. In generation 1, every allele in the “gene pool” for *D*_*O*_, *a, 𝒳* and *τ* was randomly assigned to the genomes of agents within the predator/prey populations. In each generation, the top-50 fittest agents (defined below) were paired consecutively (1^st^ and 2^nd^, 3^rd^ and 4^th^, …, 49^th^ and 50^th^), producing 4 offspring agents per pair populating the next generation. Values for offspring genes were selected uniformly randomly between the parent allele values. A Poisson process (*λ= 1*) determined the number of mutations, applied as one gene randomly altered to a new allele value for a randomly selected agent.

For each population and generation, we recorded median values of wake duration and wake bouts, and *D*_*O*_, *a, 𝓍* and *τ*. For each activity time series *R*_*g*_, order parameters were used to summarize the phenotype. We computed the complex vector:

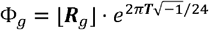

where ***T*** is time (hours). The floor of ***R***_*g*_ was taken to exclude transitions. The angle of *Φ*_*g*_ (range [0,2*ττ*)) represents average wake phase (temporal niche) and the magnitude of *Φ*_*g*_ (range [0-1]) represents strength of this tendency. Average state for each population, *Φ*_*population*_, was calculated by averaging *Φ*_*g*_ across agents.

We calculated the time series *S*_prey_ and *S*_predator_, representing the average fraction of time awake for each population. Fitness was calculated based on an agent’s overlapping wake time with each population, assuming detriment to prey during wake overlap with predators.

We define the fitness function 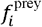 for prey agent *i*:

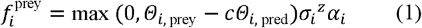

and the fitness function 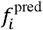 for predator agent *i*:

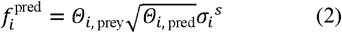

where, *Θ*_*i*, population_ *= R*_*i*_ ***S***_population_ quantifies average wake overlap of agent *i* with the specified population.

In (1), max *(0, Θ* _*i*, prey_ – *cΘ*_*i*, pred_) represents positive and negative effects of interactions with prey and predators, respectively, with floor of zero. *c* controls the detriment of prey-predator interactions due to predation and prey needing to devote resources to evade predators. *σ*_*i*_ is the fraction of time asleep, where *z* is an exponent controlling the benefit of sleep on fitness. The term *α* _*i*_ is the fraction of time awake, representing beneficial effects of waking activities, including foraging. These factors multiply as they jointly influence reproductive likelihood.

In (2), *Θ* _*i*, prey_ represents the benefit of predation opportunities. *Θ*_*i*, pred_ represents positive effects of predator-predator interactions, applying the square-root function for diminishing returns. 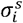 represents beneficial effects of sleep, where the exponent *s* controls the dependence on sleep duration.

For a given simulation, parameters *c, z* and *s* were kept constant (Table I). The parameter *z* was fixed at 1.6 to obtain a sleep:wake ratio close to 1:1 in the prey population; the average ratio found in phylogenetic comparative analyses of mammalian sleep [23] [24].

**TABLE 1.** Fitness function parameters.

| Physiological Model |  |
| --- | --- |
| $c$ | Predation effect on prey |
| $z$ | Modulates benefit of sleep duration for prey |
| $s$ | Modulates benefit of sleep duration for predators |
| Low-dimensional Model |  |
| $A$ | Benefit of interactions with own species |
| $b$ | Cost/benefit of predator-prey interactions |
| $c$ | Benefit of sleep duration for prey |
| $d$ | Benefit of sleep duration for predators |

In this model, we often observed coupled oscillations between populations across generations, which were stochastic due to random mutations. While oscillation frequency and phenotype varied depending on fitness parameter values, the same fundamental phenomenon was observed: over generations, prey escape to a new temporal niche, and subsequently predators invade this niche (Fig. 1). This ‘temporal niche pursuit’ resembles the classic Lotka-Volterra model, with oscillations in temporal niche rather than population size.

**FIG. 1.**
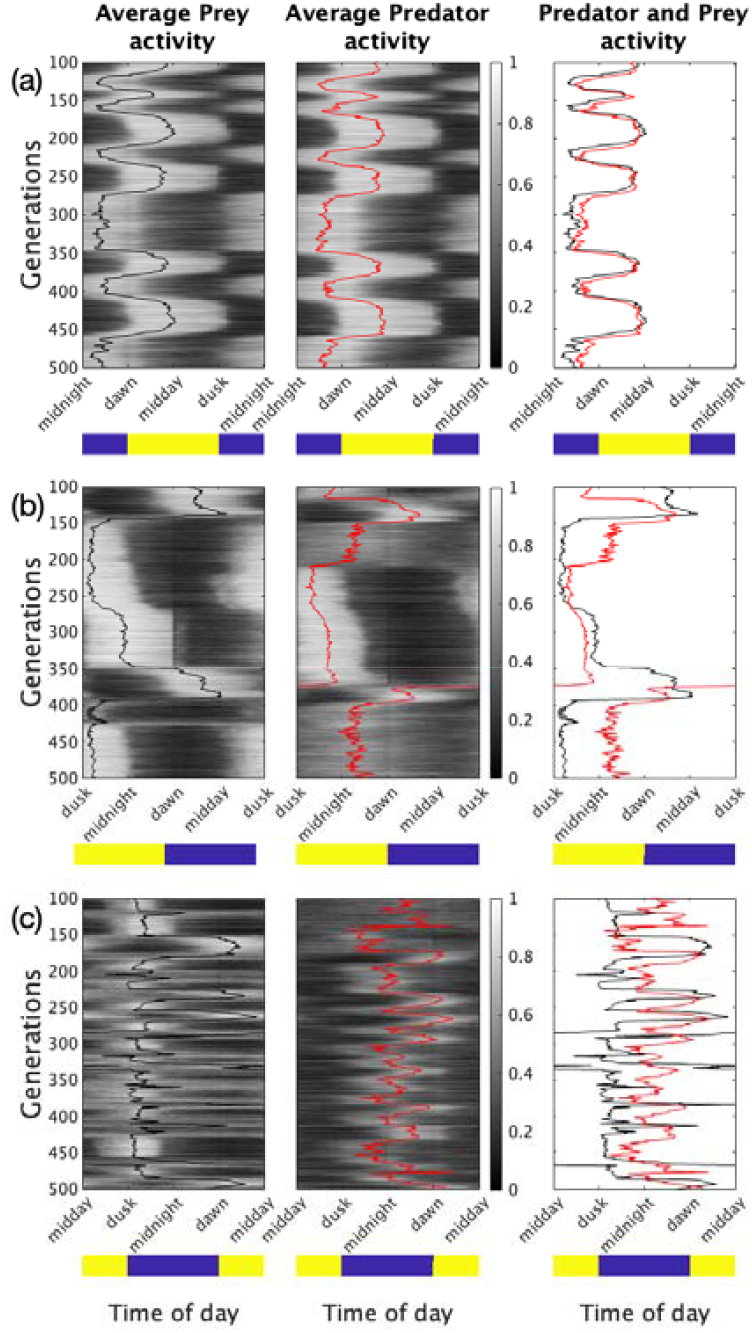
Raster plots of average prey and predator behaviors for the physiological model. Fitness parameters: (a) *c* = 0.6, *s* = 1; (b) *c* = 1, *s* = 2; and (c) *c* = 1.6, *s* = 3.5. Shaded regions indicate the proportion of population awake (black = 100% sleep, white = 100% wake). Solid lines indicate the circular average phase for prey and predators (black and red, respectively). The right column shows the two average lines overlaid. Yellow and blue bars indicate day and night, respectively.

To systematically investigate, we searched the fitness parameter space, varying *c* (0.2-3.2 in 0.2 increments) and *s* (0.5-4.0 in 0.5 increments) for oscillations. Stochastic oscillations required we develop empirical criteria for oscillation detection (Supplementary material section I). Similar maps and dynamics across random seeds (Fig. S2, S3) demonstrated robustness.

As *c* increases, predation is increasingly detrimental for prey fitness. As *s* increases, the predator’s sleep duration increases (Fig. S1), resulting in fewer predator-prey interactions. Consequently, oscillatory dynamics depend on a balance between *s* and *c*, reflected by a diagonal band of oscillatory behavior in Fig. 2(a). To the left of the band, weak incentives exist for prey to evade predators and/or predators to remain awake to prevent oscillatory dynamics. Conversely, to the right of the band, strong detrimental predation effects on prey caused rapid changes in the prey phenotypes, which were difficult for predators to consistently pursue.

**FIG. 2.**
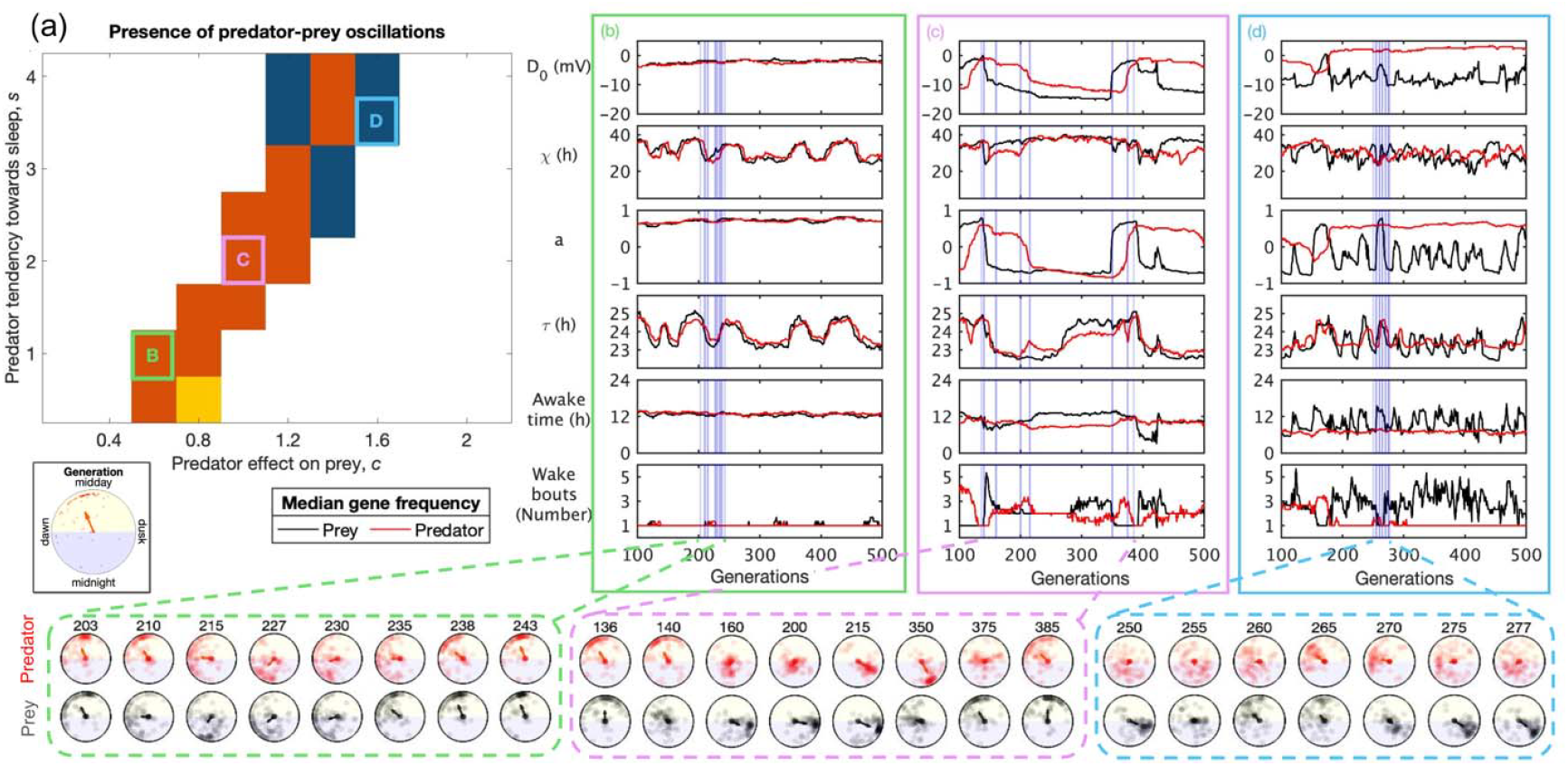
Dynamics of the physiological model. (a) Predator-prey oscillations observed across the parameter space for and . Orange = oscillations in τ and other gene(s); Blue = oscillations in only ; Yellow = oscillations in non-gene(s). Panels (b), (c), and (d) show time series across generations 100-500 (black = prey, red = predator) corresponding to labeled points in (a) for,,,, average awake time and wake bouts per day. Agent-level behaviors are shown at specific generations (blue lines) chosen from a cycle. For each generation, agents are indicated as points, while population average state,, is shown as arrows.

Oscillations were driven by different genes. Oscillations in *τ* (circadian period) were most common. In many cases, we observed coevolutionary oscillations in *τ* and another gene (e.g., *τ* and *a* in Fig. 2(b)). The higher prevalence of *τ* oscillations over other genes may be because *τ* variation enables incremental shifts in sleep timing, which can simultaneously accommodate predator avoidance and partially overlapping wake with other prey. Single point mutations can alter *τ* [25], making this a plausible mechanism for temporal niche oscillations in the evolutionary record. Although less common, we also observed oscillations in *a*, which controls diurnality vs. nocturnality (Fig. 2(b)). The relatively lower occurrence of *a* oscillations may be due to costs of evolving sleep in anti-phase with the agent’s population. Interestingly, in Fig. 2(c) we observe a case of asymmetric gene involvement where prey exhibited *τ* and *a* oscillations, whereas predators pursued prey through *τ* oscillations only. These findings demonstrate that temporal niche pursuit can arise through a variety of physiological mechanisms.

## III. LOW-DIMENSIONAL MODEL OF PREDATOR-PREY SLEEP EVOLUTION

We next investigated evolutionary dynamics using a simple, six-equation model with two populations, *x* (predator) and *Y* (prey). Each population is divided into four sleep phenotype subpopulations: nocturnal short sleep (ns), nocturnal long sleep (nl), diurnal short sleep (ds) and diurnal long sleep (dl). We assumed fixed population size for predators and prey and, under continuum limits, modeled relative fractions for each phenotype as ordinary differential equations. Fitness was defined for each phenotype, with greater reproduction for higher fitness values.

Fitness for the prey phenotype *p* at generation *t* was defined:

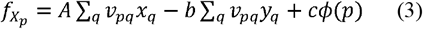

where *A> 0* represents the benefit of interactions with the same species (including reproductive opportunities); represents the cost of interactions with predators; represents the benefit of longer sleep; ; is the fraction of time that phenotypes and are mutually awake on average (see Table S2 for numerical derivation); and are the fraction of prey and predator populations in phenotype, as a function of time; and for long sleep phenotypes and otherwise.

Fitness for the predator phenotype at generation was similarly defined:

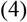

including benefit from interactions with prey; and represents the benefit of longer sleep.

The absolute fitness functions defined in (3) and (4) were scaled to relative fitness values (bounded from 0-1) using the functions

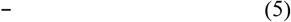

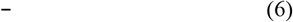

where and are the means of the four relative fitness functions for the prey and predator species, respectively.

The dynamics of the model were described by a system of ordinary differential equations, with the form

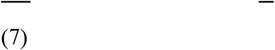

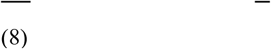

where and are the phenotypes and is the mutation rate (range 0-1). The term represents reproduction of the same phenotype with no mutation occurring, while the term represent**s** reproduction by other phenotypes that randomly mutate into phenotype (we assume mutations cause transitions from a phenotype into one of the other three possible phenotypes with equal probabilities).

The terms and are constants chosen to maintain constant population sizes defined by:

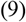

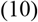

Fixed population sizes result in two conservation equations: and . Thus, the 8 phenotypes (4 prey and 4 predator) are fully described by 6 differential equations (3 prey and 3 predator).

Without loss of generality, we set in Eqn. (3, 4) and assumed (see similar results for in Fig. S4). We mapped the dynamics of this model in fitness parameter space varying the relative benefit of sleep to predators vs. prey, and, the predator-prey interaction term, with fixed . For each parameter set combination, the model was run for 101,000 generations. Each scenario was simulated from 50 initial conditions, drawn randomly from the interval [0,1] obeying the conservation equations above. A fast Fourier transform was applied to the last 100,000 generations for each predator and prey phenotype. The dominant period is plotted in Fig. 3(a), requiring minimum power of 1 for detection, and using the median dominant period across the 50 initial conditions.

**FIG. 3.**
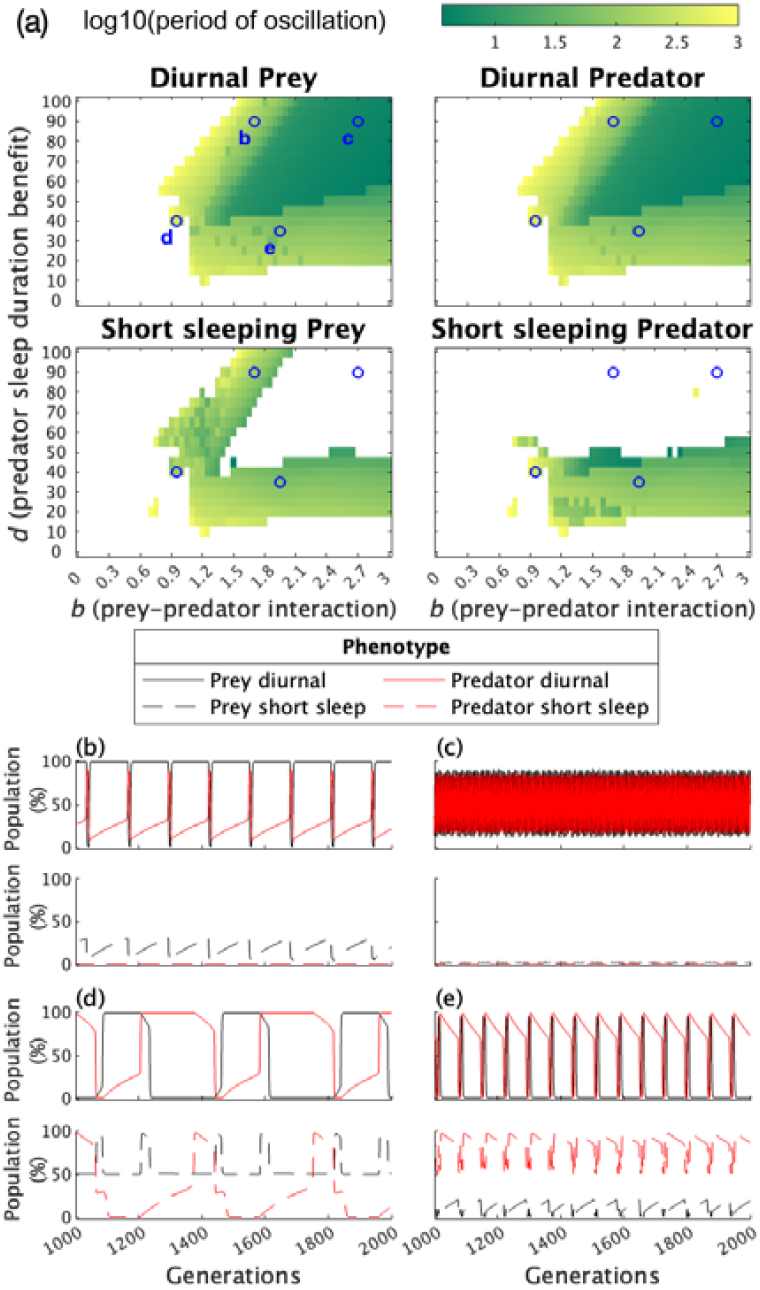
Low-dimensional model of predator-prey sleep evolution. (a) Map of detectable oscillations in each phenotype (diurnal vs. nocturnal; short vs. long sleep) for prey and predators in fitness parameter space. Panels (b-e) show characteristic cases of phenotype time series across generations 1000-2000 (black = prey, red = predator) corresponding to the points labeled in panel (a).

This low-dimensional model recapitulated many characteristics of the physiological model. First, the diagonal band of stable oscillations in fitness parameter space (Fig. 3(a)) resembled the diagonal band in the physiological model, again arising fro**m** interplay between the strength of predator-prey interactions and the sleep benefit to predators. Second, multiple modes of oscillatory dynamics emerged, echoing the physiological model (Fig. 3(b-e)). Where there was strong incentive for predators to sleep and prey to evade predators, both populations maintained long-sleep but oscillated rapidly in their temporal niche phenotype (period of ≲10 generations) (Fig 3(c)). Conversely, a weaker prey-predator interaction (Fig. 3(b)) resulted in longer-period oscillations (∼100-1000 generations), where both populations experienced temporal niche oscillations, but prey additionally experienced sleep duration oscillations. A lower predator sleep bia**s** (Fig. 3(d), 3(e)) resulted in mixed-mode oscillations for both populations. Lower predator sleep bias and weaker predator-prey interaction caused slower oscillations and were in some cases double-peaked (e.g., prey sleep duration in Fig 3(d)).

## IV. CONCLUSIONS

By applying a physiologically derived model to investigate sleep evolution, we uncovered ‘temporal niche pursuit’: a phenomenon in which prey recurrently escape predator interactions by seeking **a** distinct temporal niche, and predators pursue prey by invading this niche. We found multiple modes of genetic variation could drive these oscillatory dynamics. This was observed in both the physiological model and a low-dimensional model, suggesting this is a fundamental outcome for coevolution of sleep patterns in a predator-prey system.

Of the few attempts to model sleep evolution [26], only one is a dynamical predator-prey agent-based model [27],[28]. This model demonstrated prey avoidance of predators but the potential for long-term dynamics was not investigated, and it was not based on physiology. In our models, we assume fixed-sized populations and non-overlapping generations. With one prey population, we implicitly assumed a specialist predator. Different assumptions could result in other dynamics [29]. We simulated a constant light-dark cycle and excluded other factors that could affect activity, such as environmental temperature. Furthermore, we focused on sleep physiology, whereas temporal niche shifts may occur concomitantly with other adaptations, including thermoregulation and eye structure accommodating differing light levels. While such adaptations could be rate-limiting, there is empirical evidence that eye structure can rapidly change to accommodate changes in temporal niche, though photoreceptor adaptations may occur more slowly [30], [7].

Temporal niche varying continually over evolutionary timescales may explain phenomena such as stark differences in temporal niche between some closely related mammalian species [11] and high diversity in temporal niche within species [31]. For some species, niche switching can even occur within individual animals resulting from environmental conditions [3,32,33] or predation pressure [34], suggesting temporal niche is a flexible evolutionary trait [26]. The existence of temporal niche pursuit reconciles the famous ‘nocturnal-bottleneck hypothesis’ with more recent evidence of dynamic changes in temporal niche among both the early mammals and contemporaneous dinosaurs.

This project was supported by Australian Research Council projects DP210102924 and DP220102812. We are thankful for use of MonARCH (Monash Advanced Research Computing Hybrid). AJKP has received funding from Versalux and Delos, and he is co-founder of Circadian Health Innovations Pty Ltd.

## REFERENCE

[1] H. J. Kanaya et al., A sleep-like state in *Hydra* unravels conserved sleep mechanisms during the evolutionary development of the central nervous system, Sci. Adv. 6, eabb9415 (2020).

[2] R. D. Nath, C. N. Bedbrook, M. J. Abrams, T. Basinger, J. S. Bois, D. A. Prober, P. W. Sternberg, V. Gradinaru, and L. Goentoro, The jellyfish *Cassiopea* exhibits a sleep-like state, Curr. Biol. 27, 2984 (2017).

[3] R. A. Hut, N. Kronfeld-Schor, V. van der Vinne, and H. De la Iglesia, In search of a temporal niche: environmental factors, Prog. Brain Res. 199, 281 (2012).

[4] K. O. Lear, N. M. Whitney, J. J. Morris, and A. C. Gleiss, Temporal niche partitioning as a novel mechanism promoting co-existence of sympatric predators in marine systems, Proc. Biol. Sci. 288, 20210816 (2021).

[5] I. Castro-Arellano and T. E. Lacher, Temporal niche segregation in two rodent assemblages of subtropical Mexico, J. Trop. Ecol. 25, 593 (2009).

[6] M. Albrecht and N. J. Gotelli, Spatial and temporal niche partitioning in grassland ants, Oecologia 126, 134 (2001).

[7] M. P. Gerkema, W. I. L. Davies, R. G. Foster, M. Menaker, and R. A. Hut, The nocturnal bottleneck and the evolution of activity patterns in mammals, Proc. Biol. Sci. 280, 20130508 (2013).

[8] R. Refinetti, The diversity of temporal niches in mammals, Biol. Rhythm Res. 39, 173 (2008).

[9] G. S. van Doorn, J. Schepers, R. A. Hut, and A. T. Groot, Sex-specific expression of circadian rhythms enables allochronic speciation, Evol. Lett. 9, 65 (2025).

[10] A. W. Crompton, C. R. Taylor, and J. A. Jagger, Evolution of homeothermy in mammals, Nature 272, 333 (1978).

[11] R. Maor, T. Dayan, H. Ferguson-Gow, and K. E. Jones, Temporal niche expansion in mammals from a nocturnal ancestor after dinosaur extinction, Nat Ecol Evol 1, 1889 (2017).

[12] L. Schmitz and R. Motani, Nocturnality in dinosaurs inferred from scleral ring and orbit morphology, Science 332, 705 (2011).

[13] R. F. Nespolo, L. D. Bacigalupe, C. C. Figueroa, P. Koteja, and J. C. Opazo, Using new tools to solve an old problem: the evolution of endothermy in vertebrates, Trends Ecol. Evol. 26, 414 (2011).

[14] J. P. Hayes and T. Garland, The evolution of endothermy: testing the aerobic capacity model, Evolution 49, 836 (1995).

[15] J. Wiemann, I. Menéndez, J. M. Crawford, M. Fabbri, J. A. Gauthier, P. M. Hull, M. A. Norell, and D. E. G. Briggs, Fossil biomolecules reveal an avian metabolism in the ancestral dinosaur, Nature 606, 522 (2022).

[16] K. D. Angielczyk and L. Schmitz, Nocturnality in synapsids predates the origin of mammals by over 100 million years, Proc. Biol. Sci. 281, (2014).

[17] P. A. Robinson, A. J. K. Phillips, B. D. Fulcher, M. Puckeridge, and J. A. Roberts, Quantitative modelling of sleep dynamics, Philos. Trans. A Math. Phys. Eng. Sci. 369, 3840 (2011).

[18] S. Postnova, A. Layden, P. A. Robinson, A. J. K. Phillips, and R. G. Abeysuriya, Exploring sleepiness and entrainment on permanent shift schedules in a physiologically based model, J. Biol. Rhythms 27, 91 (2012).

[19] A. J. K. Phillips, B. D. Fulcher, P. A. Robinson, and E. B. Klerman, Mammalian rest/activity patterns explained by physiologically based modeling, PLoS Comput. Biol. 9, e1003213 (2013).

[20] A. J. K. Phillips, P. A. Robinson, D. J. Kedziora, and R. G. Abeysuriya, Mammalian sleep dynamics: how diverse features arise from a common physiological framework, PLoS Comput. Biol. 6, e1000826 (2010).

[21] A. C. Skeldon, D.-J. Dijk, and G. Derks, Mathematical models for sleep-wake dynamics: comparison of the two-process model and a mutual inhibition neuronal model, PLoS One 9, e103877 (2014).

[22] M. Magnin, M. Rey, H. Bastuji, P. Guillemant, F. Mauguière, and L. Garcia-Larrea, Thalamic deactivation at sleep onset precedes that of the cerebral cortex in humans, Proc. Natl. Acad. Sci. U. S. A. 107, 3829 (2010).

[23] J. A. Lesku, T. C. Roth, N. C. Rattenborg, C. J. Amlaner, and S. L. Lima, Phylogenetics and the correlates of mammalian sleep: a reappraisal, Sleep Med. Rev. 12, 229 (2008).

[24] P. McNamara, I. Capellini, E. Harris, C. L. Nunn, R. A. Barton, and B. Preston, The phylogeny of sleep database: A new resource for sleep scientists, Open Sleep J. 1, 11 (2008).

[25] M. Menaker and J. S. Takahashi, Genetic analysis of the circadian system of mammals: properties and prospects, Seminars in Neuroscience 7, 61 (1995).

[26] S. L. Lima and N. C. Rattenborg, A behavioural shutdown can make sleeping safer: a strategic perspective on the function of sleep, Anim. Behav. 74, 189 (2007).

[27] A. Acerbi and C. L. Nunn, Predation and the phasing of sleep: an evolutionary individual-based model, Anim. Behav. 81, 801 (2011).

[28] C. L. Nunn, D. R. Samson, and A. D. Krystal, Shining evolutionary light on human sleep and sleep disorders, Evol Med Public Health 2016, 227 (2016).

[29] U. Dieckmann, P. Marrow, and R. Law, Evolutionary cycling in predator-prey interactions: population dynamics and the red queen, J. Theor. Biol. 176, 91 (1995).

[30] J. Baker and C. Venditti, Rapid change in mammalian eye shape is explained by activity pattern, Curr. Biol. 29, 1082 (2019).

[31] R. Costa-Pereira, V. H. W. Rudolf, F. L. Souza, and M. S. Araújo, Drivers of individual niche variation in coexisting species, J. Anim. Ecol. 87, 1452 (2018).

[32] N. Mrosovsky and S. Hattar, Diurnal mice (Mus musculus) and other examples of temporal niche switching, J. Comp. Physiol. A Neuroethol. Sens. Neural Behav. Physiol. 191, 1011 (2005).

[33] J. G. Davimes, A. N. Alagaili, M. F. Bertelsen, O. B. Mohammed, J. Hemingway, N. C. Bennett, P. R. Manger, and N. Gravett, Temporal niche switching in Arabian oryx (*Oryx leucoryx*): Seasonal plasticity of 24 h activity patterns in a large desert mammal, Physiol. Behav. 177, 148 (2017).

[34] M. G. P. Fenn and D. W. Macdonald, Use of middens by red foxes: risk reverses rhythms of rats, J. Mammal. 76, 130 (1995).

