## Supplementary information for "Temporal niche pursuit in a simulated evolution of sleep/wake patterns"

### I. DEFINING OSCILLATIONS IN "GENES" IN THE PHYSIOLOGICAL MODEL

Since the oscillations observed were highly stochastic, we developed a set of empirical criteria for oscillation detection described in this section. This oscillation detection method was used in all physiological model simulations, including in supplementary section IV, where different seeds were trialled.

First, a smoothed correlation vector was generated:

1. Log transformation of $\chi$ gene: For the $\chi$ gene only, a natural log transformation of the data was taken.

2. Detrending median gene data: These median values of gene frequency were linearly detrended across the 500 generations to create stationary time series. This involved taking the linear regression of the raw data and subtracting the average slope from the raw data. This was done so that correlation values calculated subsequently did not include slow moving genetic drifts over the timescale of greater than hundreds of generations, as we were strictly looking for oscillations (with transitions happening over a timescale of fewer than ~50 generations).

3. Calculating standard deviation of median gene data: The standard deviation of the stationary median gene data for both predator and prey genes were recorded. A greater value for standard deviation would indicate a greater variability in the data and hence a greater likelihood of oscillations being found. This was used in condition 5.

4. Smoothed correlation vector: The cross-correlation between the detrended median gene values were computed for lag values of 0-100 generations, to identify lagged cycles (i.e., predators pursuing prey). For example, a lag of 1 between the predator and prey median tau values would be the median tau prey values from generation 1 to 499 and median predator tau values from generation 2 to 500. A lag of 30 would be the median tau prey values from generation 1 to 470 and median predator tau values from 31 to 500. We would expect a larger lag would mean a lower correlation between the two (predator and prey) time series. If oscillations were to occur (and present in both populations), we'd expect that a peak in the correlation value to occur at a lag value greater than 1.

   The lag vector (0:100) thus has an associated "correlation" vector of equal length (101) which gives a numerical indication of how closely a change in the median gene in the prey population is associated with a change in the median gene of the predator population over increasingly widening generation windows. The cross-correlation function was smoothed with a 5-point moving mean to reduce the effect of noise. This smoothed correlation vector was used in conditions 1, 2, and 4.

For a particular set of fitness parameter values to be said to contain genetic oscillations across generations, the smoothed cross-correlation function was required to satisfy all 5 of the following conditions for a particular gene.

**Condition 1:** The maximum value in smoothed correlation vector must be greater than 0.4, indicating the presence of strongly correlated prey-predator behavior.
**Condition 2:** The smoothed correlation vector had to have at least one peak or local maximum.
**Condition 3:** The maximum value of non-smoothed correlation vector must not occur at lag = 0. A maximum correlation value occurring at lag 0 would indicate that there was no "pursuit" of the prey by the predator population.
**Condition 4:** The range in the smoothed correlation values must be greater than 0.55, indicating a distinct oscillatory frequency.
**Condition 5:** Finally, to identify which specific gene(s) were undergoing oscillations, we required a meaningful amount of genetic variation across the oscillation. This was indicated by the standard deviation of the median gene value, calculated across generations, exceeding a threshold percentage of the allowed range for that gene. Since model dynamics have different sensitivity to each gene, we specified thresholds for each parameter, such that coupled oscillations between predator and prey populations could be isolated. Thresholds for $D_0$, $a$, $\chi$ and $\tau$ were 6.5%, 12.0%, 5.3%, and 8.5%, respectively.
While for every gene, conditions 1-4 were constant, in condition 5, the standard deviation threshold was varied depending on the gene. The standard deviation of the median value of the gene in both the predator and prey populations had to be met for the condition to be satisfied.

The percentage of standard deviation calculated for the predators and prey was taken as a fraction of the range of possible gene values [max-min value given in main text, repeated here: (i) offset in the sleep drive, $D_0$ (-30 to 13 mV), (ii) the sleep homeostatic time constant, $\chi$ (5 to 45 h, (iii) the diurnality index, $\alpha$ (-1 to 1), and (iv) the intrinsic circadian period, $\tau$ (22 to 26 h)]. This standard deviation percentage had to meet the specified thresholds above, indicating a minimum amount of variability in the predator and prey median gene values independent of their interactions with the other population.

### II. DEFAULT PARAMETER VALUES USED IN THE HARD-SWITCH MODEL

TABLE S1: Fixed parameter values in physiological model.

| Neural population parameters | Default value |
|---|---|
| $\nu_{vd}$ (mV s) | -0.17 |
| $\nu_{vh}$ (mV s) | 1 |
| $A_0$ (mV) | 1.3 |
| **Homeostatic parameters** | |
| $\mu$ (mV) | 10 |
| **Circadian modulation parameters** | |
| $\nu_{\mathrm{md}}$ (mV) | 0.01 |
| k ($s^{-1}$) | 17 |
| $\delta$ | 2.8 |
| $b$ ($s^{-1}$) | 4.8 |
| **Masking by light parameters** | |
| $\nu_{\mathrm{vb}}$ | 0 |
| **Circadian parameter values** | |
| $\gamma$ | 0.13 |
| $c1$ | 1/3 |
| $c2$ | 4/3 |
| $c3$ | 0 |
| $c4$ | -256/105 |
| $q$ | 1/3 |
| $h$ | -0.2 |
| **Photic drive parameters** | |
| $\beta$ ($h^{-1}$) | 0.4 |
| $\alpha_0/I_0$ | $0.1/9500^{0.5}$ |
| G' | 37 |
| $p$ | 0.5 |
| $I_1$(lux) | 100 |
| $r$ | 0.4 |

### III. AVERAGE PREY AND PREDATOR SLEEP DURATIONS IN PHYSIOLOGICAL MODEL

For the simulations presented in the main paper, we also computed the average prey and predator sleep duration values across the parameter space for $c$ and $s$. The results are shown in Fig. S1.

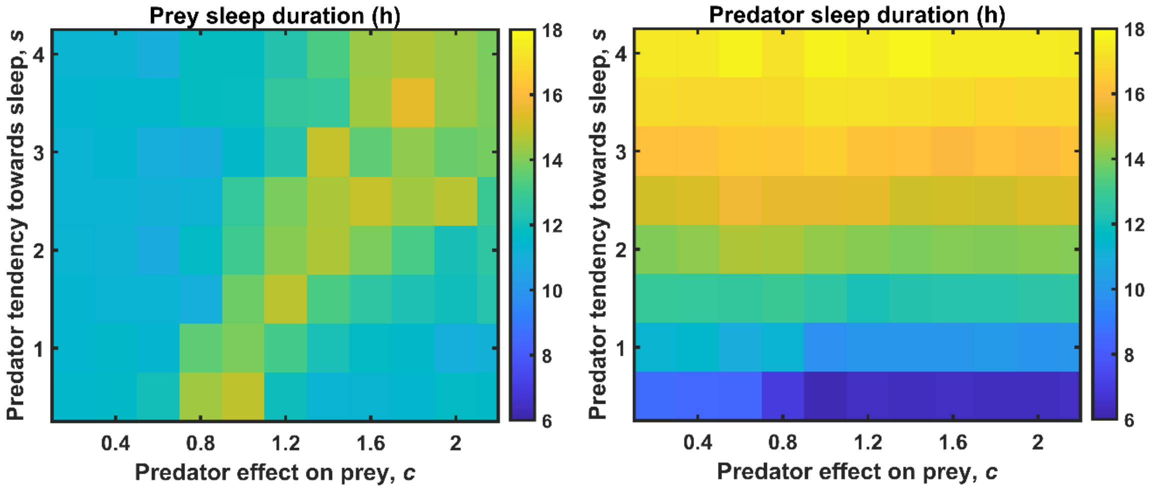


FIG. S1. Average prey and predator sleep durations for the physiological model, corresponding to Fig. 3.

### IV. TRIALLING DIFFERENT RANDOM SEEDS IN PHYSIOLOGICAL MODEL

Simulations were performed with different random seed values to test robustness of the solutions, shown in Fig. S2. The default value for the seed value is 1 if no seed value was explicitly specified in MATLAB. Results throughout the paper are for seed value 1.

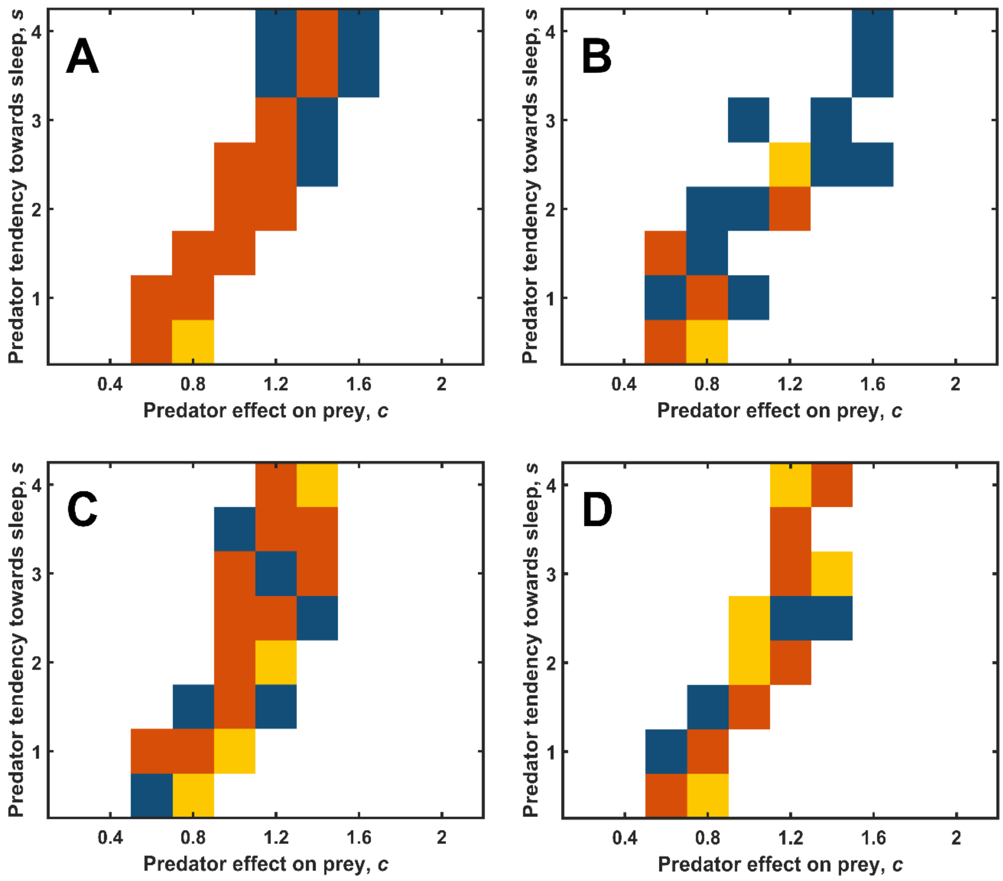

FIG. S2. The replicability of the oscillations is tested with different seeds. The oscillations with the same set of coefficients are given. The same set of conditions outlined in SECTION I of the supplementary material are used to determine if oscillations occur in particular genes.

### IV. TRIALLING DIFFERENT RANDOM SEEDS IN PHYSIOLOGICAL MODEL (CONTINUED)

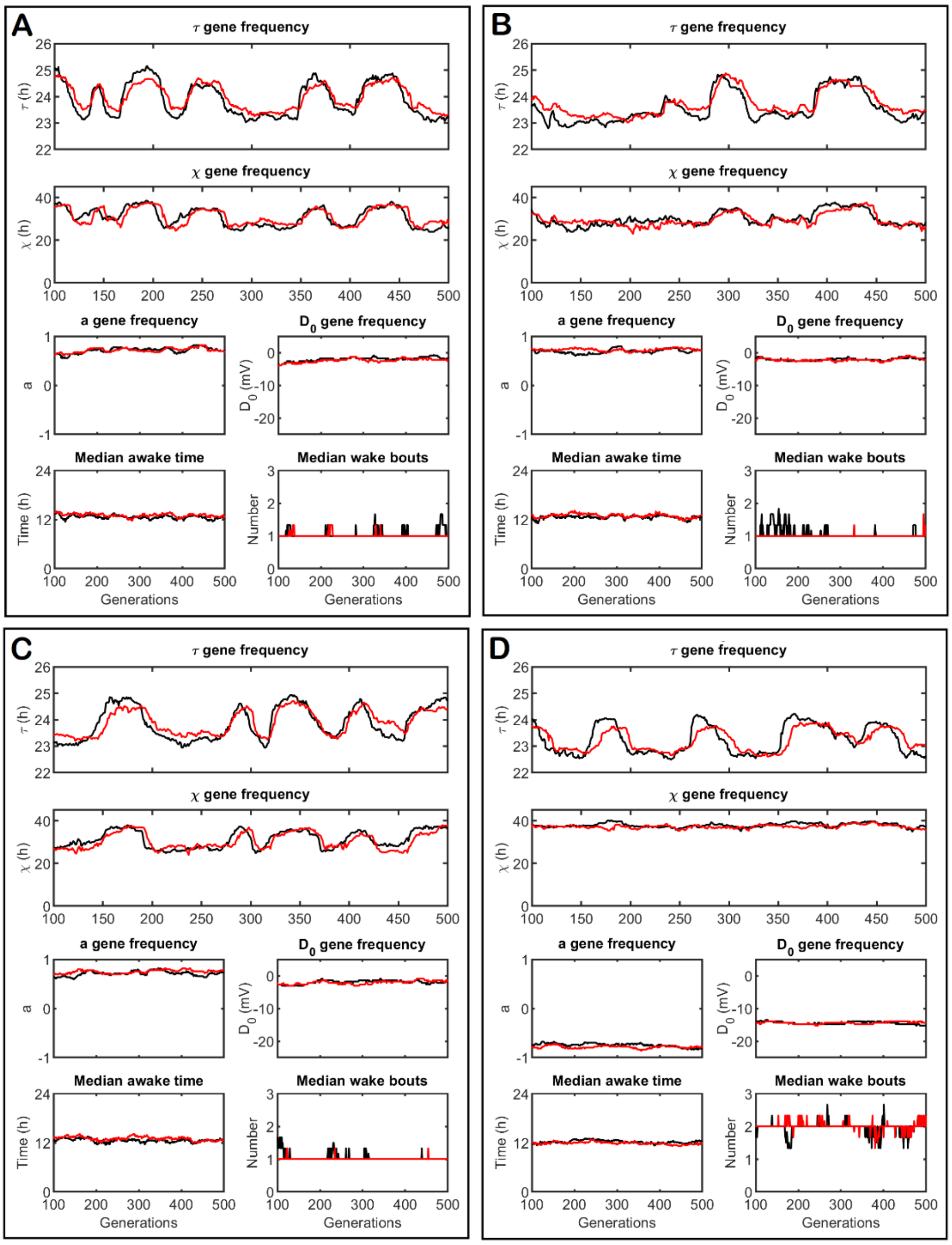


FIG. S3. Outputs from the physiological model simulated for 4 different random seeds, using the parameter values c= 0.6, s = 1, and z = 1.6, for four different seed values (1, 2, 3, 4 for A, B, C, D).

A run keeping the coefficient set constant (parameter set 26 – [0.6, 1.6, 1]) with seeds incremented from 1 to 100. Across these 100 runs, 96 exhibited $\tau$ oscillations, 28 exhibited $\chi$ oscillations, and no other genes exhibited oscillations (according to the conditions outlined above). If, for a particular seed, $\chi$ gene oscillations occurred, they occurred alongside $\tau$ gene oscillations.

### V. ASSUMED SLEEP/WAKE PATTERNS IN THE SIMPLIFIED MODEL

We simulated sleep/wake patterns for the four phenotypes in the simplified model. The day was divided into four equal sections of 6-hour duration. The timing of these four sections is arbitrary. We assumed 6-hour sleep durations for the short sleep phenotypes and 12-hour sleep durations for the long sleep phenotypes. Crosses in Table S2 represent sleep periods for each phenotype. The amount of overlapping wake time per day between two phenotypes then corresponded to the number of columns where neither phenotype had a cross (e.g., 2 columns = 12 hours for the phenotypes ds and dl).

TABLE S2: Assumed sleep/wake patterns for the four phenotypes in the simplified model (diurnal short sleep [ds], diurnal long sleep [dl], nocturnal short sleep [ns], nocturnal long sleep [nl]).

| | 9pm-3am | 3am-9am | 9am-3pm | 3pm-9pm |
|---|---|---|---|---|
| ds | X | | | |
| dl | X | X | | |
| ns | | | X | |
| nl | | | X | X |

### VI. SIMPLIFIED MODEL DYNAMICS WITH MUTATION RATE = 0.1

Finally, we tested the model dynamics using a higher mutation rate (0.1) to determine how this affected the periodic dynamics in the model. The results are displayed in Fig. S4.

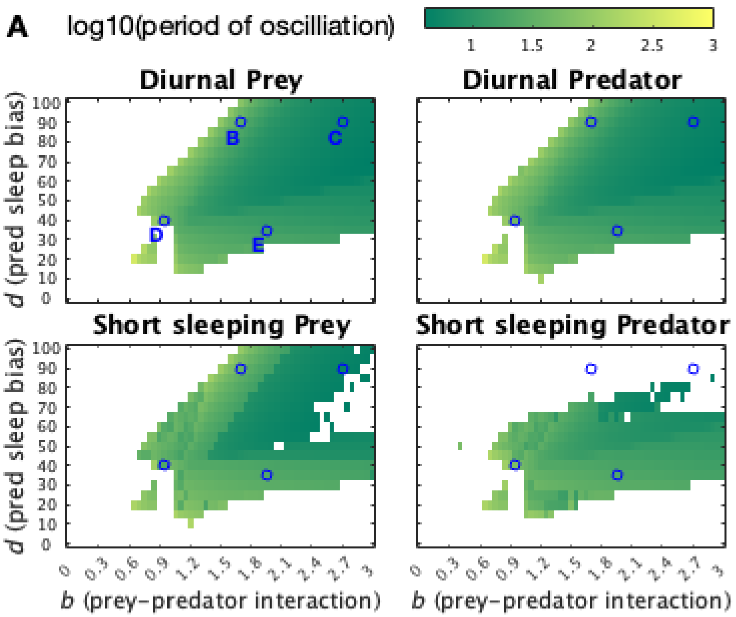


FIG. S4. Dynamics of the simplified model with mutation rate set to 0.1. Legend is as per Fig. 3A.
